# The acquisition of additional feedback loops may optimize and speed up the response of quorum sensing

**DOI:** 10.1101/2021.06.11.448020

**Authors:** Marco Fondi, Francesca Di Patti, Elena Perrin

## Abstract

Bacterial quorum sensing (QS) is a cell-to-cell communication system in which specific signals are activated to coordinate, for example, pathogenic behaviors and help bacteria collectively respond to perturbations. QS in Gram-negative bacteria is typically regulated by a N-acyl-homoserine lactone (AHL) molecules-mediated system, homologous of *Vibrio fischeri* LuxI-R. In many cases, bacteria possess more than one QS system, based on different types of molecules, that interact through a complex regulatory network. Presumably, these configurations have emerged over time from simpler ones through the acquisition of novel players (e.g. transcription factors) that have been successfully integrated into the native regulatory systems. However, the advantages provided by these alternative/additional configurations on QS-related phenotypes is poorly predictable only based on their underlying network structure. Here, we have adopted a modelling approach to infer the possible improvements conferred by the acquisition of additional control over bacterial regulation of QS. We use the *Burkholderia* genus as a case study because some of these strains, besides the LuxIR-like system (named CepIR), possess an integrated regulatory module named CciIR that interferes with the CepIR system through the implementation of several positive and negative control loops. Being associated to a genomic island (cci island), this additional module is prone to being horizontally transferred, giving rise to a potentially patchy genomic distribution and, in turn, to a *complete* (CepIR and CciIR systems together) vs. core (CepIR only) organization of QS regulation in this group of microorganisms. By using both deterministic and stochastic modelling we show that, upon their activation, the two regulatory schemes may lead to different phenotypes and to distinct responses to the extracellular concentration of signalling molecules. In particular, our simulations show that the presence of the additional regulatory module may confer specific improvements, including a faster response time and optimized control of QS regulation. Interestingly, some of these features may be particularly advantageous during host invasion, thus highlighting once more the importance of QS in the establishment and maintenance of bacterial infections.

## 2. Introduction

Quorum sensing (QS) is a communication tool among bacterial cells that allows to coordinate collective behaviours depending on cell density and on the composition of the surrounding microbial community [1, 33]. It is based i) on communication molecules of different chemical nature, called autoinducers (AI), that are produced and secreted by bacterial cells and ii) on receptor regulators that, after the recognition of a given amount of a specific AI, can control the expression of several different genes [50]. In Gram-negative bacteria QS is typically regulated by N-acyl-homoserine lactone (AHL) molecules-mediated system. This is the case of the model organism *V. fischeri* where LuxI synthesizes an AHL signal and LuxR is an AHL receptor protein that activates or represses gene expression by binding to a consensus sequence in the promoter regions of target genes [48]. Many different cellular processes are under the control of QS, including virulence, pathogenesis, biofilm formation and fluorescence [36]. They are generally highly expensive from an energetic point of view, and bacterial cells activate them only when a large population can benefit from them. As a consequence, the regulation of this phenotype is tightly controlled by microorganisms that, in time, have evolved many different regulatory strategies. These, in turn, may give origin to complex behaviours that include life-style switches whose evolutionary benefit resides in the capacity of controlling and reacting to environmental changes [17]. In many cases, bacteria possess more than one QS system, possibly based on different type of molecules, that interact through a complex regulatory network that can also respond to inducers produced by other bacteria [50, 36, 20]. A paradigmatic example of this complex regulation is represented by the *Burkholderia cepacia* complex (Bcc), a group of more than 20 bacterial species, with an evolving taxonomy, characterized by a large variety of different lifestyles [9, 47]. The LuxIR homolog CepIR was the first QS system to be identified in a *B. cenocepacia* strain [21]. Afterwards, some *B. cenocepacia* representatives have been shown to possess two main pairs of QS regulation modules: i) the CepIR system which is present in all species of the Bcc [14, 3, 21] and ii) the CciIR system, whose distribution is much more limited and that is thought to be present only in *B. cenocepacia* strains containing the cenocepacia island (cci) found in the highly transmissible ET12 strains [4]. N-octanoyl-homoserine lactone (C8-HSL) and of N-hexanoyl-homoserine lactone (C6-HSL) are known to be the main AIs for QS regulation in *Burkholderia* [21]. Concerning their production, CepI directs the synthesis of C8-HSL and minor amounts of C6-HSL whereas the AHL synthase CciI mostly produces C6-HSL and minor amounts of C8-HSL. Further, the two regulators of these systems (CepR and CciR) define a hierarchical regulatory relationship (Figure 1) where CepR, besides negatively regulating its own expression, is required for *cepI* and the *cciIR* operon expression, while CciR negatively regulates its own expression, as well as that of *cepI* [4].

**Figure 1:**
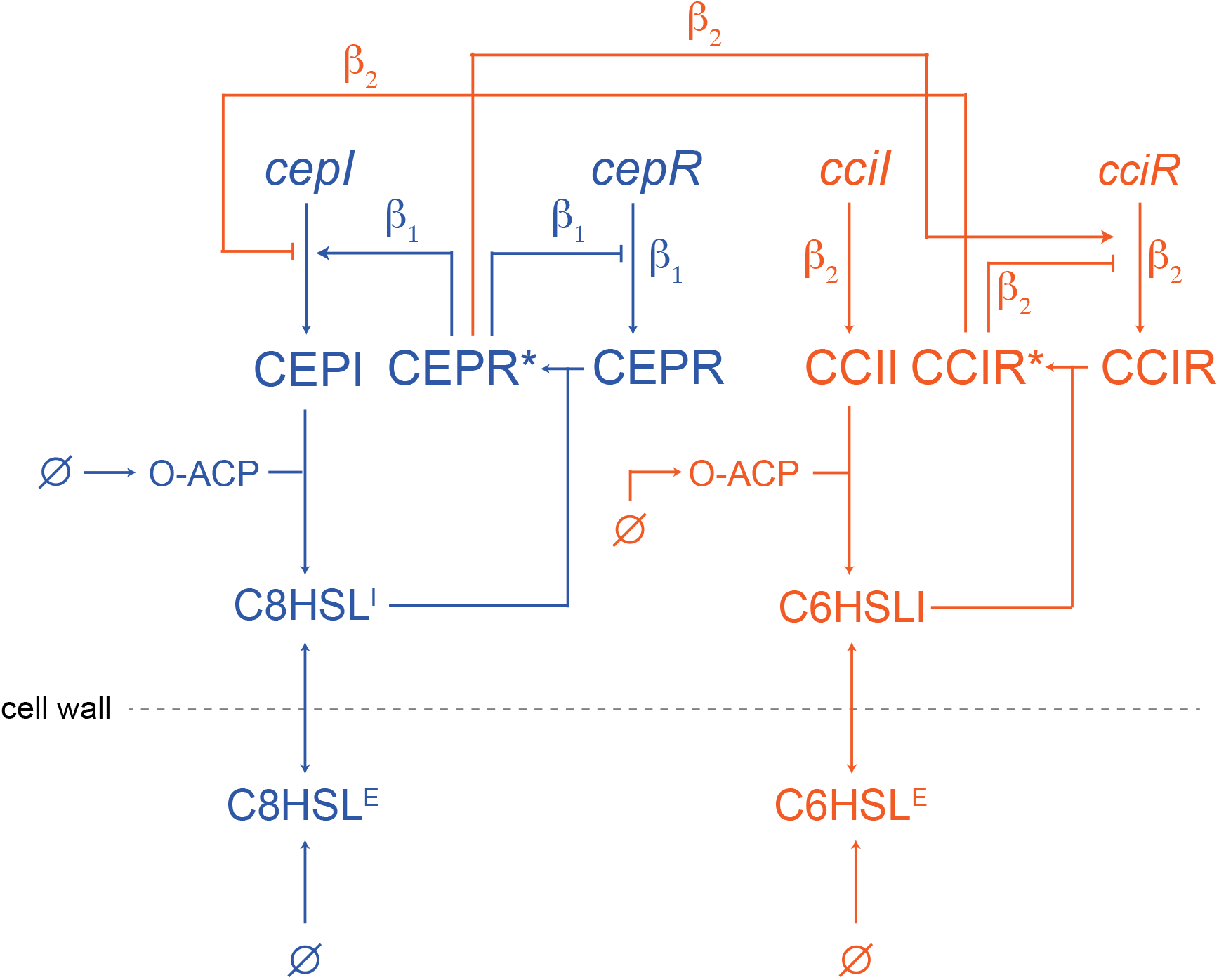
Schematic representation of the complete *Burkholderia cenocepacia* QS regulatory network modelled in this work (full model). The part of the model colored in blue represents the *core* model. The blue part plus the orange part represent the *complete* model. *β*_1_ and *β*_2_ indicate the basal transcription rates (*β*) and were both set to *β* for simulating the *complete* architecture. *β*_2_ was set to 0 when simulating the *core* architecture.

The inclusion of a regulatory system (CciIR) inside a genomic island (*cci*) represents an extraordinary example of the plasticity of the cellular transcriptional network and an exceptional occasion to study the effect of alternative regulatory architectures on the expression of the resulting phenotypic trait. Indeed, horizontal gene transfer (HGT) and/or recombination events can lead to the appearance of novel regulatory modules (here, CciIR) that, to be maintained over evolutionary time scales, have to integrate with the pre-existing one(s) (here, CepIR) and, eventually, increase the fitness of the recipient organism [5]. This is of particular importance in the context of the genus *Burkholderia* since its representatives can colonize different environments, including the respiratory tract of immunocompromised patients, like people with Cystic Fibrosis (pwCF) [8, 49, 34]. Although the incidence of infection by Bcc species in pwCF is not high in comparison with other CF pathogens, the mortality rate is high [11, 16], due to a combination between the high virulence of these species and their high levels of intrinsic AR and of QS-related traits in general [41]. Thus, understanding the intimate functioning of their QS regulation may provide additional hints on the emergence of phenotypic traits that are tightly linked to the harmfulness of this group of bacteria.

Here, we have used comparative genomics to determine the exact distribution of the *cci* island, and in particular of *cciI* and *cciR*, inside the *Burkholderia* genus. Given that the presence/absence pattern of these genes pointed towards the occurrence of several gene gain/loss events in the evolution of the genus, we next asked which may be the advantages conferred by the acquisition of the CciIR regulatory system and its integration in one of the native QS regulation system. To answer this question we have built a comprehensive dynamic model that is capable of simulating the regulation of QS for those strains possessing a *complete* (i.e. CepIR and CciIR) or a *core* (only CepIR) architecture. Thus, from a more general perspective, we have modelled the consequences of evolution-driven changes in the structure of a regulatory circuit on the overall dynamics of cellular behaviour.

Our simulations suggest that the implementation of additional positive and negative feedback loops may confer specific improvements to the *complete* QS-regulation system, including a faster response time and an overall optimized control of QS regulation in response to the concentration of extracellular AI. Interestingly, some of these features appear to depend on cellular growth rate, with a response optimum identified in a specific range of possible growth rates and matching the one that is most likely found during bacterial infections.

## 3. Results

### 3.1. The distribution of cci genes in the Burkholderia genus is compatible with extensive gene gain/loss

Despite part of the *cci* genomic island is being used as an epidemic strain marker [4], its actual distribution inside *Burkholderia* has not been investigated at the genus level. For this reason, we probed the presence of the *cci* island encoded genes in all the available complete *Burkholderia* genomes and combined this information with the genomic relatedness of the representatives of this genus. This latter metrics was computed by calculating the Average Nucleotide Identity (ANI, see Methods) for each pair of *Burkholderia* genomes included in the dataset. As reported in Supplementary Information (Figure S1), the distribution of the genes by the *cci* island follows a pattern that matches the one obtained through ANI computation. More in detail, in the cluster comprising *B. mallei, B. pseudomallei* and *B. thalilandiensis* species, the complete set of *cci* genes is never found. Instead, the distribution of *cci* genes in the other cluster (hereinafter cci-group, that includes, among the others, the BCC and the members of the *B. glumae/gladioli/plantarii* group) is patchy and includes (22) microbes that harbour more than 50% of the reference *cci* genes and others (93) possessing less than 50% of the reference *cci* genes (Figure 2). This pattern suggests a complex evolutionary history for this genomic islands, mainly guided by gain/loss events. Despite the detailed reconstruction of the steps that led to the actual distribution of the *cci* island in *Burkholderia* is beyond the scope of this work, two main scenarios may explain it. Either the *cci* island was acquired once in the ancestor of the extant *cci*-group and then lost in some of its representatives or, alternatively, it was acquired independently multiple times in the evolution of this group. In either of these scenarios, to be maintained over evolutionary time, *cci* genes have had to successfully integrate into the regulatory network of their hosts and to eventually increase their fitness. Although in some cases the contribution of newly acquired genes to the overall cellular fitness is easily explainable (e.g. antimicrobial resistance, virulence, etc.), in some others identifying the advantages they confer is not straightforward. This might be the case, for example, of additional, horizontally transferred, transcription factors that may interact with the native regulatory network in unpredictable ways and giving rise to complex phenotypes. Here, we focus our attention on two genes included in the *cci* genomic island, namely *cciI* and *cciR*. Both of them are known i) to be involved in QS regulation and ii) to be perfectly integrated in the native QS regulation system encoded by CepI and CepR [32] (Figure 1). To be maintained over evolutionary time, this new regulatory system should provide some (currently undisclosed) improvements to the strains harbouring it. More explicitly, here we ask which may be the consequences and advantages (if any) provided by the acquisition of a novel, additional QS control system and by its integration into the native regulatory circuit?

**Figure 2:**
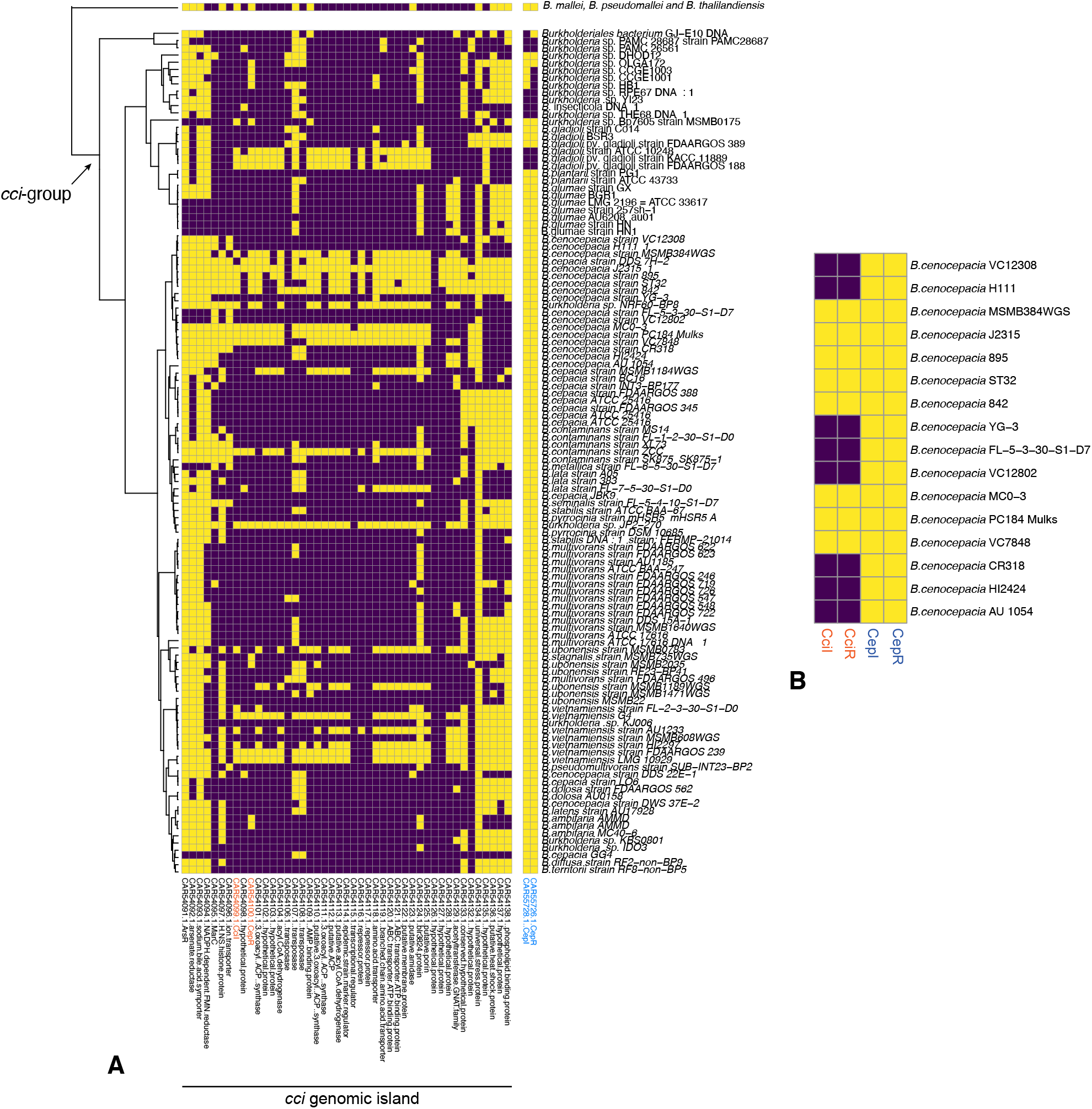
A) Distribution of *cci* encoded genes in a selected set of *Burkholderia* representatives. Yellow and purple squares indicate the presence or the absence of the corresponding gene, respectively. QS regulatory systems are coloured in orange (CciIR) and blue (CepIR). B) Focus on CepIR and CciIR regulatory systems in some *B. cenocepacia* representatives.

### 3.2. A mathematical model for the core and complete QS regulation systems

The distribution of *cciI, cciR, cepI* and *cepR* genes inside *Burkholderia* (Figure 2A) points towards the presence of two (alternative) QS regulatory circuits within the *B. cenocepacia* species (and *Burkholderia* in general) (Figure 1): i) a *core* regulatory circuit, composed by CepI, CepR and CepR* (the active form of CepR), that is peculiar of those organisms missing *cciI* and *cciR* and ii) a *complete* system (resulting from the integration of CepIR and CciIR) characteristic of those *Burkholderia* representatives harbouring the four genes (Figure 2B).

It is important to stress that, in some organisms, this regulatory scheme might actually include other genes and/or TFs [33]. This is the case, for example, of *B. cenocepacia* J2315 in which there exists a third LuxR homolog (CepR2) that lacks an associated AHL synthase and that is negatively regulated by CciR [26].

However, since i) these genes, in most cases, do not influence the expression of the other two main systems (CepIR and CciIR) modelled herein and, more importantly, ii) we do not have enough detailed information on the kinetics of these additional regulators, they were not included in the mathematical formulation of QS regulation presented in this work.

The QS model implemented in this work is schematically represented in Figure 1. The model takes into account 4 genes (*cepI, cepR, cciI* and *cciR*), 4 proteins (CepI, CepR, CciI and CciR), 2 activated transcription factors (CepR* and CciR*) and 3 metabolites (C8HSL, C6HSL, O-ACP, octanoyl acyl carrier protein). The latter represents the most likely substrate for the production of AI molecules [30]. As for metabolites, OACP is only present inside the cell whereas we define two versions of both C8HSL and C6HSL. These include the intracellular (C8HSl^I^ and C6HSl^I^) and extracellular (C8HSl^E^ and C6HSl^E^) counterparts. This allows modelling the presence of the autoinducer synthesized by other cells to the extracellular space (production of C8HSl^E^ and C6HSl^E^) and its uptake into the cytoplasm (conversion of C8HSL^E^ / C6HSL^E^ to C8HSl^I^ / C6HSl^I^). Endogenously produced C8HSl^I^ and C6HSl^I^ can either be secreted (i.e. converted back to C8HSL^E^ and C6HSL^E^) or remain inside the cell and bind to CepR and CciR and convert them to their active form (CepR* and CciR*, Figure 1). The CepIR system relies on the N-octanoyl-homoserine lactone (C8HSL) synthase CepI and the transcriptional regulator CepR that specifically binds to C8HSL becoming active. In its activated form (CepR*), CepR binds upstream to CepI and promotes its transcription. Within the circuits considered in this work, CepR* also promotes the transcription of *cciR* and represses its own transcription (Figure 1). Finally, it binds to the *cep* boxes, located upstream specific genes, causing the induction or repression of their expression [14, 24, 43]. This overall organization is mirrored by the CciIR system in which CciI synthesizes C6-HSL that can bind to CciR and convert it to the active form (CiiR*) [25]. The CciIR system also influences the CepIR system, as CciR negatively regulates *cepI* [25]. Finally, CciR* represses the transcription of *cciR*. The model further takes into account protein and metabolites degradation rates and the overall dilution effect due to cellular growth (*δ*). The entire model (hereinafter termed *complete*) recapitulates the architecture of the *Burkholderia* representatives possessing both the CepIR and CciIR regulation system (Figure 1), whereas the architecture of those species harbouring only the CepIR regulation is accounted for by the *core* model (the blue part of Figure 1). The *complete* model contains 24 parameters (Table1) and 13 of them are shared with the model accounting for the *core* architecture. We describe the *complete* model using the following chemical equations (the *core* model is basically a subset of the *complete* model and is described in Supplementary Material S1):

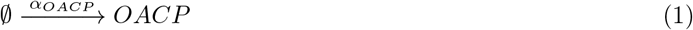

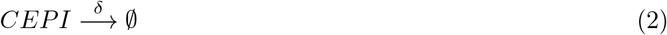

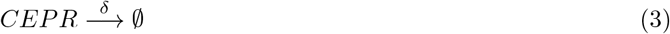

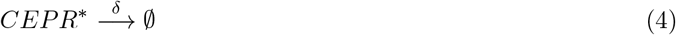

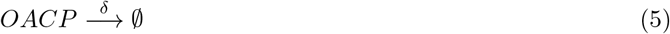

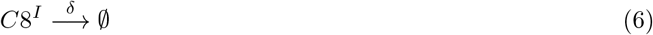

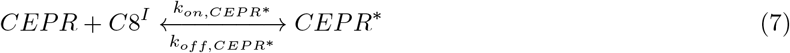

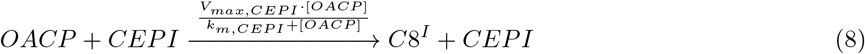

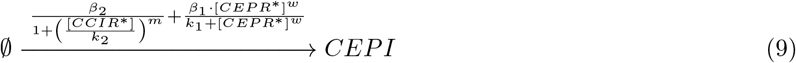

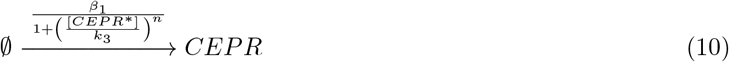

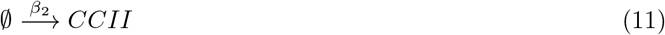

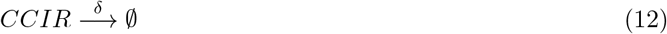

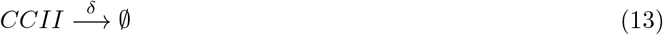

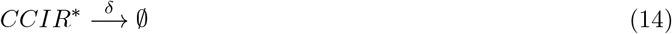

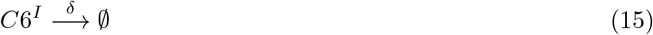

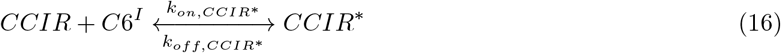

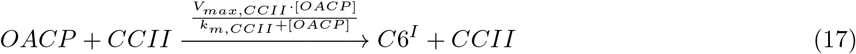

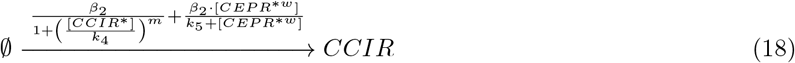

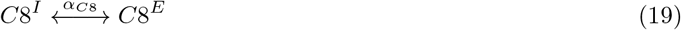

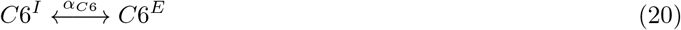

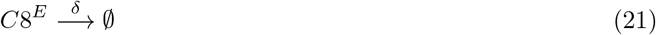

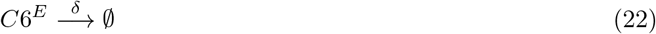

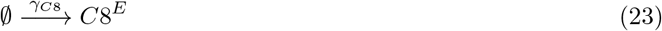

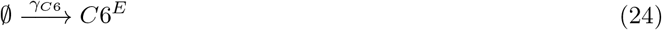

**Table 1:** Description of the parameters used in the model

| Parameter | Description | Value | Unit | Reference |
| --- | --- | --- | --- | --- |
| $\beta_1, \beta_2$ | Basic transcription rate for models M1 and M2 | 1/80 | nM min<sup>-1</sup> | Semsey et al. 2012 |
| $\delta$ | Combined protein and metabolite dilution rate | 0.001 | min<sup>-1</sup> | Boada et al. 2016 |
| $k_{on,CEPR^*}$ | $CEPR^*$ C8HSL association constant | 0.1 | min<sup>-1</sup> nM<sup>-1</sup> | Weber and Buceta 2013 |
| $k_{off,CEPR^*}$ | $CEPR^*$ C8HSL dissociation constant | 10 | min<sup>-1</sup> | Weber and Buceta 2013 |
| $\alpha_{C8^E}$ | C8HSL diffusion rate across the membrane | 10 | min<sup>-1</sup> | Weber and Buceta 2013 |
| $\alpha_{C6^E}$ | C8HSL diffusion rate across the membrane | 10 | min<sup>-1</sup> | Weber and Buceta 2013 |
| $\alpha_{OACP}$ | OACP production rate | 0.1 | min<sup>-1</sup> | Fekete et al. 2010 |
| $k_{on,CCIR^*}$ | $CCIR^*$ C6HSL association constant | 0.1 | min<sup>-1</sup> nM<sup>-1</sup> | Weber and Buceta 2013 |
| $k_{off,CCIR^*}$ | $CCIR^*$ C6HSL dissociation constant | 10 | min<sup>-1</sup> | Weber and Buceta 2013 |
| $V_{max,CEPI}$ | CEPI maximal rate | 0.0041 | min<sup>-1</sup> | Buroni et al. 2018 |
| $k_{m,CEPI}$ | CEPI Michaelis-Menten constant | $0.068 \cdot 10^{-6}$ | nM | Buroni et al. 2018 |
| $V_{max,CCII}$ | CCII maximal rate | 0.0041 | min<sup>-1</sup> | Buroni et al. 2018 |
| $k_{m,CCII}$ | CCII Michaelis-Menten constant | $0.068 \cdot 10^{-6}$ | nM | Buroni et al. 2018 |
| $\gamma_{C6/C8,colony}$ | C8/C6 diffusion from other cells | .015 | min<sup>-1</sup> | Weber and Buceta 2013 |
| $w, n, m$ | Hill's coefficients | 1.7 | — | Weber and Buceta 2013 |
| $k_1$ | $CEPR^*$ - CEPI activation coefficient | 10 | nM | Assumed |
| $k_2$ | $CCIR^*$ - CEPI/R repression coefficient | 10 | nM | Assumed |
| $k_3$ | $CEPR^*$ - CEPR repression coefficient | 10 | nM | Assumed |
| $k_4$ | $CCIR^*$ - CCII/R repression coefficient | 10 | nM | Assumed |
| $k_5$ | $CEPR^*$ - CCIR repression coefficient | 10 | nM | Assumed |

From this set of chemical equations we can derive the corresponding system of ordinary differential equations [46], describing the change in time of the concentration of each species. Defining the concentration as the ratio of the average number of molecules of each species to the cell volume *V*, and denoting such variables by the squared brackets, the dynamics of the system is given by:

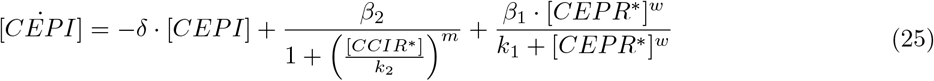

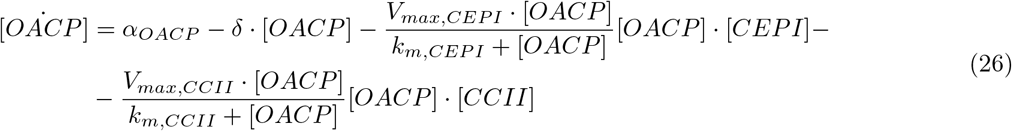

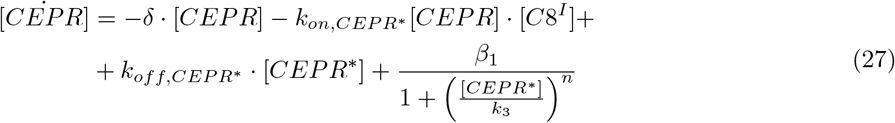

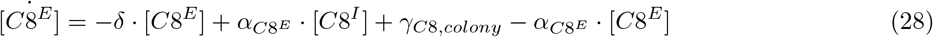

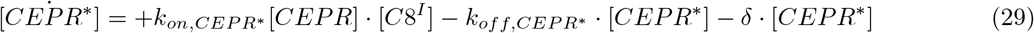

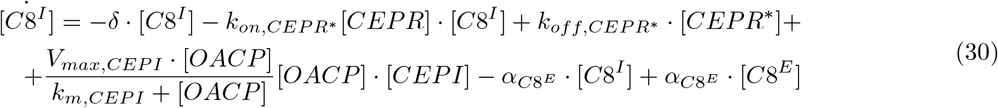

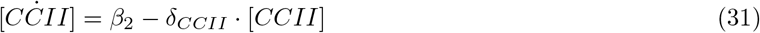

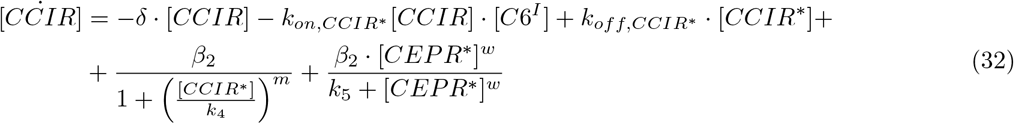

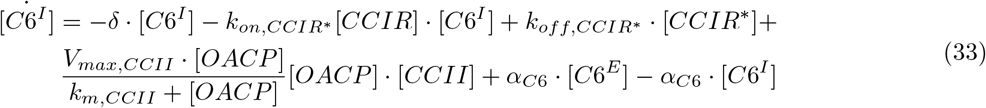

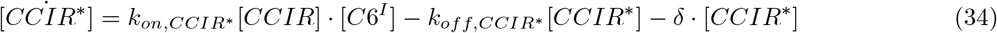

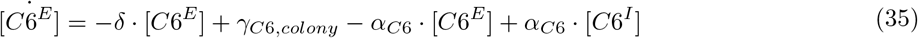

We would like to point out that we used to distinct transcription rate constants for the two modules (*β*_1_ and *β*_2_) to be able to switch on/off the CciIR system and thus to convert the *complete* model into the *core* one, simply by setting *β*_2_ =0.

### 3.3. CciIR integration with the native CepIR system may speed up the response time of QS regulation

To explore the effects of the *cciIR* system on the overall dynamics of QS regulation, we simulated the behaviour of the system for 1200 minutes upon its activation and evaluated the dynamics of the main species included in the model (Figure 3). We found that, with the exception of OACP, all the main species included in the model reach higher steady-state concentration in the *complete* model in respect to the *core* one. This is even more evident for CepI, whose steady-state concentration is particularly higher in the *complete* model. Notably the steady-state concentration of the AI (C8HSL) falls well in the range of the one experimentally measured in *B. cenocepacia* [22].

**Figure 3:**
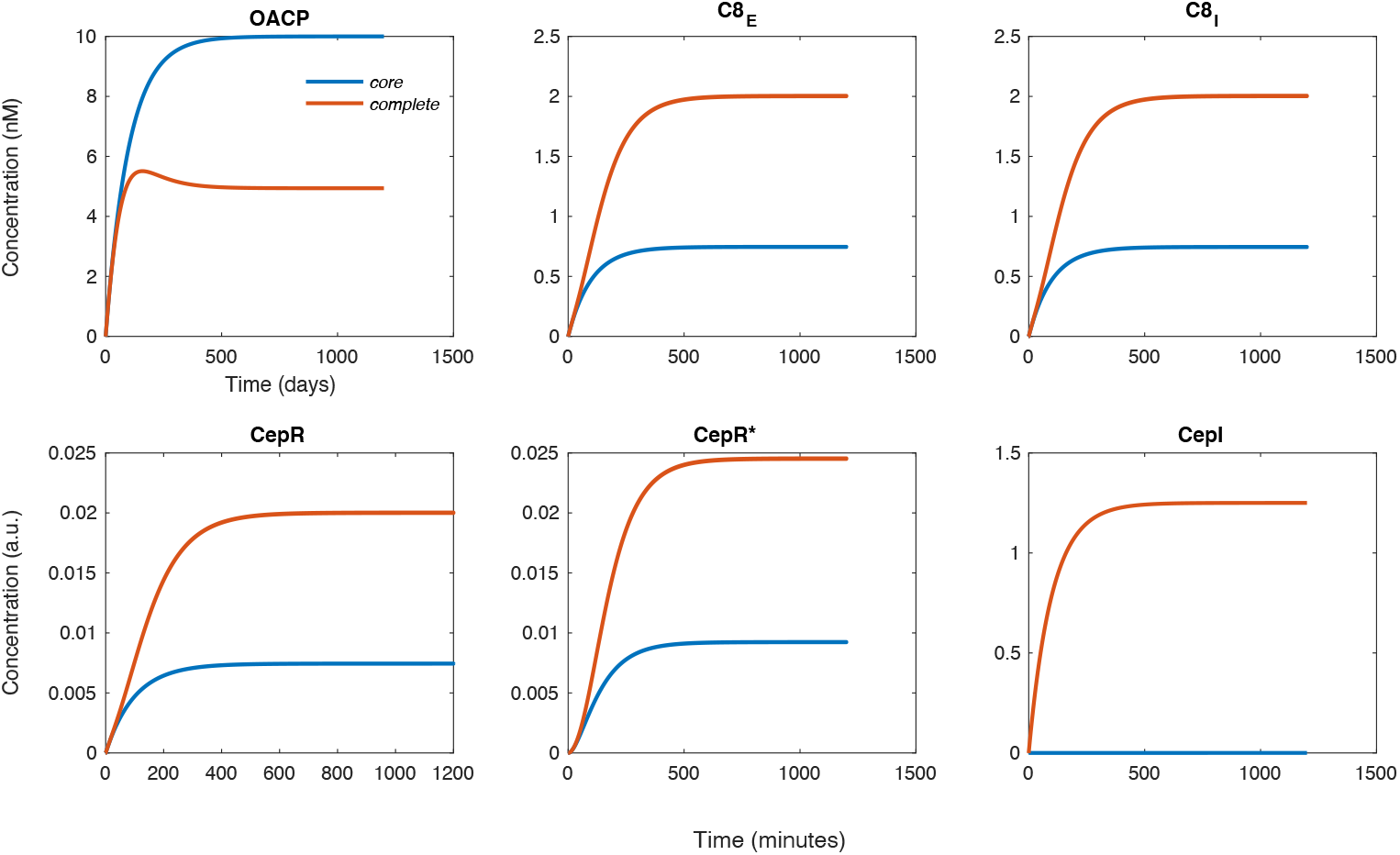
Full dynamics for all the main species included in the *core* and *complete* models

Besides steady-state concentration, regulatory circuits as the ones implemented in this work can be characterized by their response time (RT), i.e. the time required by each species to reach its half-maximal steady-state concentration. Thus, we evaluated the RT of each main player involved in QS regulation shared by the two circuits (CepI, CepR and CepR*). As shown in Figure 4, CepR maintains the very same dynamics both in the *complete* and in the *core* models. Almost the same is true for CepR*, whose RT is slightly slower in the *complete* model in respect to the *core* one. On the contrary, CepI displays a much shorter RTs in the *complete* model. According to these results, the complete model seems to guarantee a faster activation of the QS-mediated response in relation to external signals. More precisely, the RTs are 69 and 264 minutes in the case of the *complete* and *the* core models, respectively. It is to be noticed that these results were obtained using a single set of parameters (i.e. the one reported in Table 1), only a fraction of which were retrieved from *Burkholderia-specific* literature sources. For this reason we decided to inspect the robustness of the model(s) to changes in each of these parameters. Specifically, we performed a local sensitivity analysis (see Methods) and quantified how the local variation of each parameter influenced the outcome of the model. As shown in Figure 5 and 6, the model is robust to variations in the values of most parameters as local sensitivity is mostly maintained in the range of 1 * 10^−2^/1 * 10^−4^. Exceptions are represented by *α_OACP_, β_1_, γ_*C*8,colony_* and *δ* for the *core* model and *V_max,CEPI_, α_OACP_, β_1_, β_2_, δ, γ_*C*8,colony_, γ_*C*6,colony_* and *V_max,CciI_* for the *complete* model. In particular, the parameter accounting for the overall cell growht rate (*δ*) is the one that seems to affect the most the dynamics of the species in the model. This prompted us to evaluate the effect of this parameter on the two main features that seem to distinguish the two architectures, i.e. RT and the steady state values of the main species. We thus repeated the simulations described above for different values of *δ*. Concerning the response time of CepI, we found a strong influence of *δ* on the RT of the *complete* and *core* models Figure 7A. More in detail, defining ΔRT as the variation of the difference in the RT between the *complete* and the *core* circuits, we found a negative correlation between ΔRT and the simulated growth rate. In other words, as higher values of ΔRT indicate a shorter RT of the *complete* vs. the *core* model, our simulations suggest that the slower the growth rate, the faster is the RT in the *complete* model in respect to the *core* one. We also computed ΔRT for the AI molecules (C8HSL and C6-HSL) and found that, also in this case, ΔRT gets higher (i.e. the *complete* circuit is faster) at lower growth rates, with an overall trend that resembles the one found for CepI (Figure 7B). Indeed, the time required to reach its half-maximal concentration remains comparable between the two architectures between 0.1 and 0.005 h^−1^) and then gets larger (i.e. faster in the *complete* model) matching the dynamics observed for *cepI* for lower values of *δ*. Interestingly, *δ* seems to affect the steady-state concentration of C8HSL, with the complete model showing to be able to produce (and hence secrete) a quantity of C8HSL up to four time higher in respect to the *core* one, in a specific growth rate range (Figure 8).

**Figure 4:**
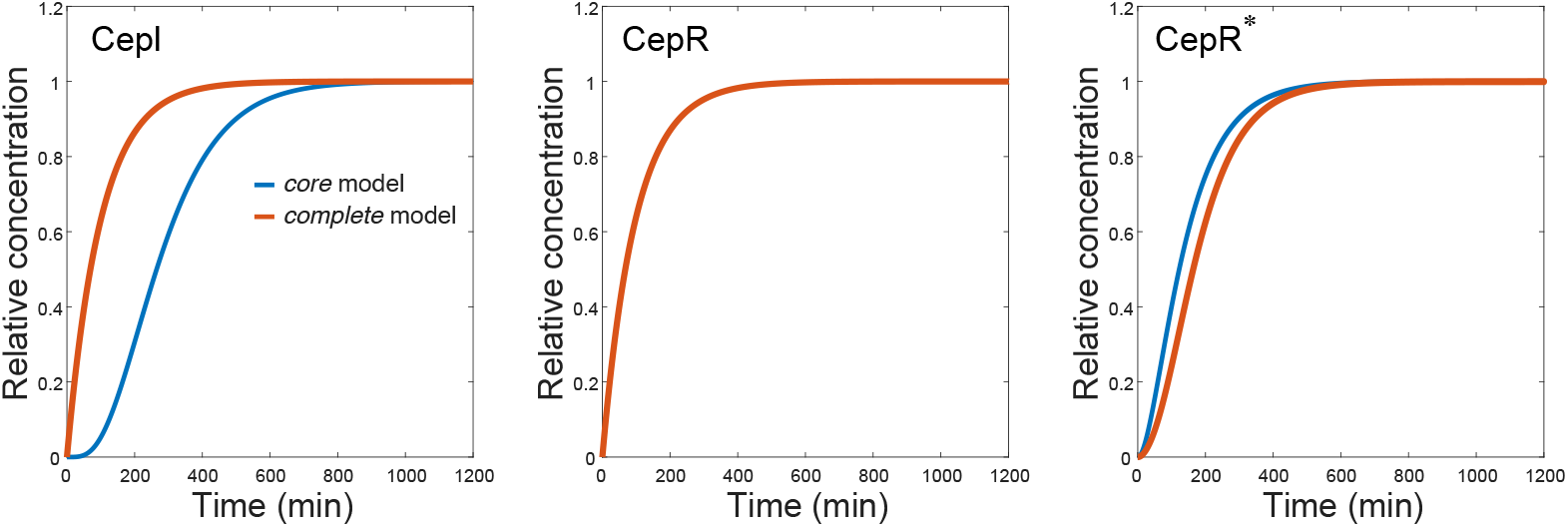
Dynamics of CepI, CepR and CepR* in the complete (orange) and core (blue) models

**Figure 5:**
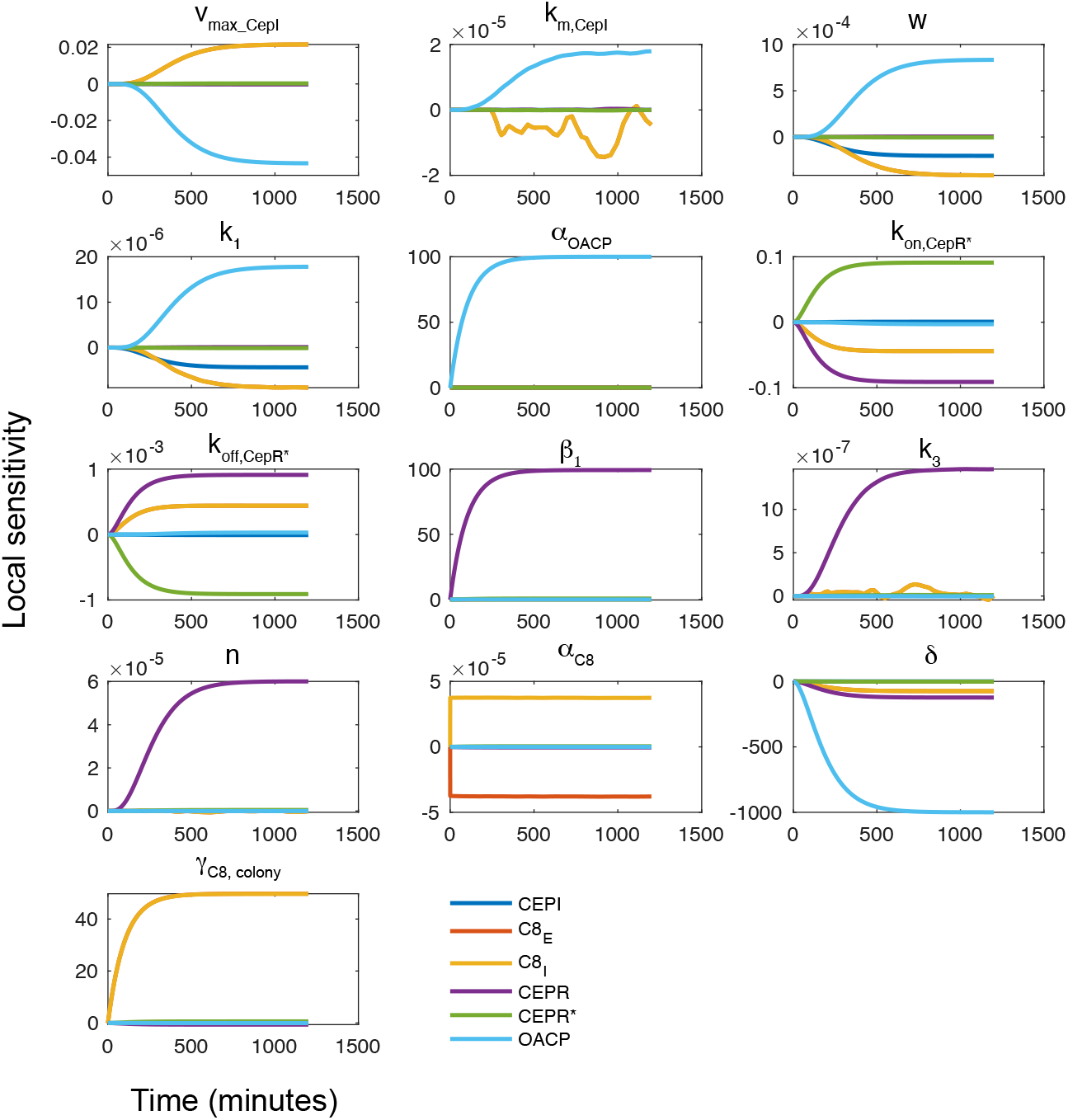
Local sensitivity analysis for the *core* model. Each plot represents a sensitivity analysis for the parameter specified above the plot. Each coloured line is representative of each of the remaining parameters in the model.

**Figure 6:**
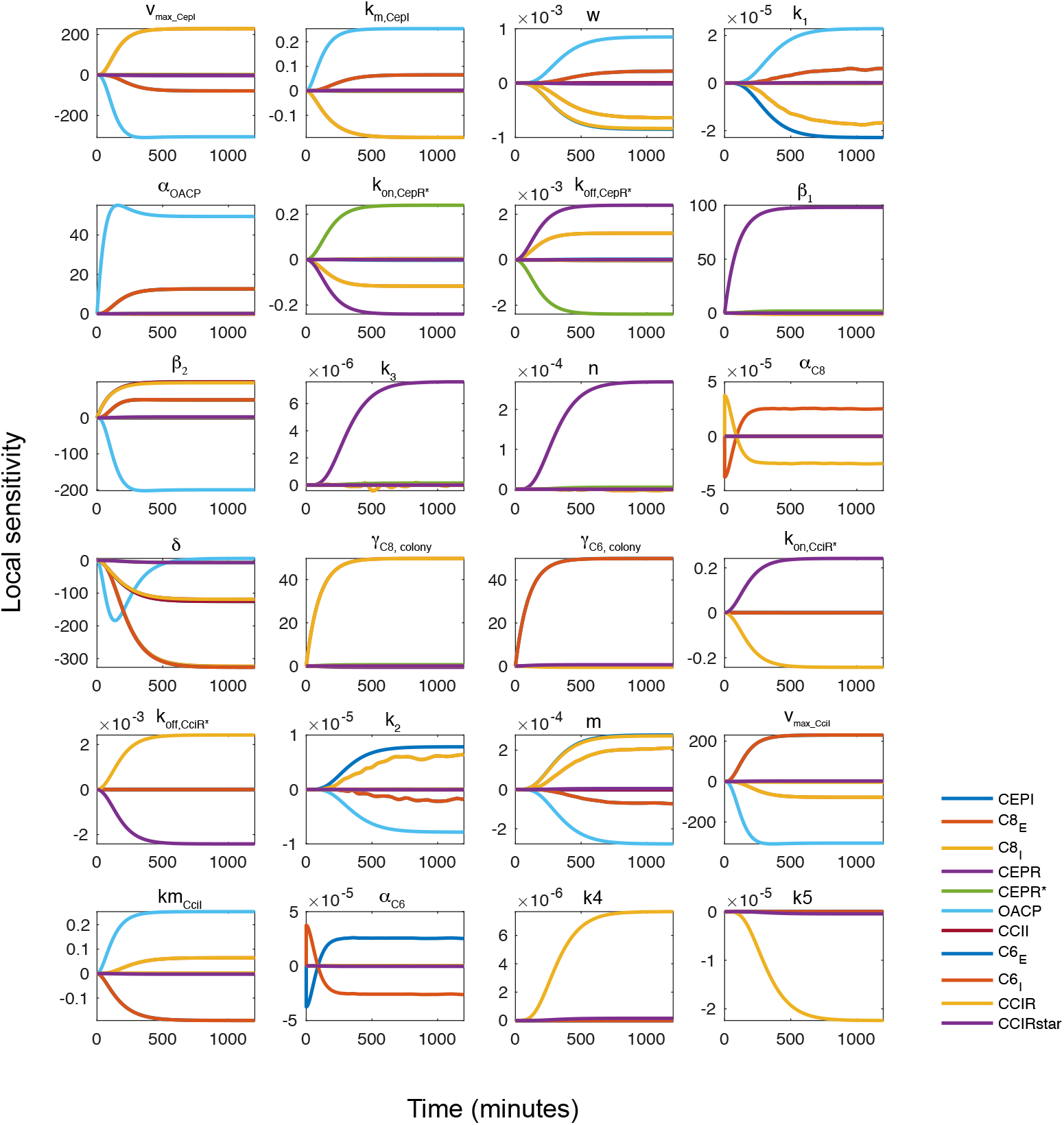
Local sensitivity analysis for the *complete* model. Each plot represents a sensitivity analysis for the parameter specified above the plot. Each coloured line is representative of each of the remaining parameters in the model.

**Figure 7:**
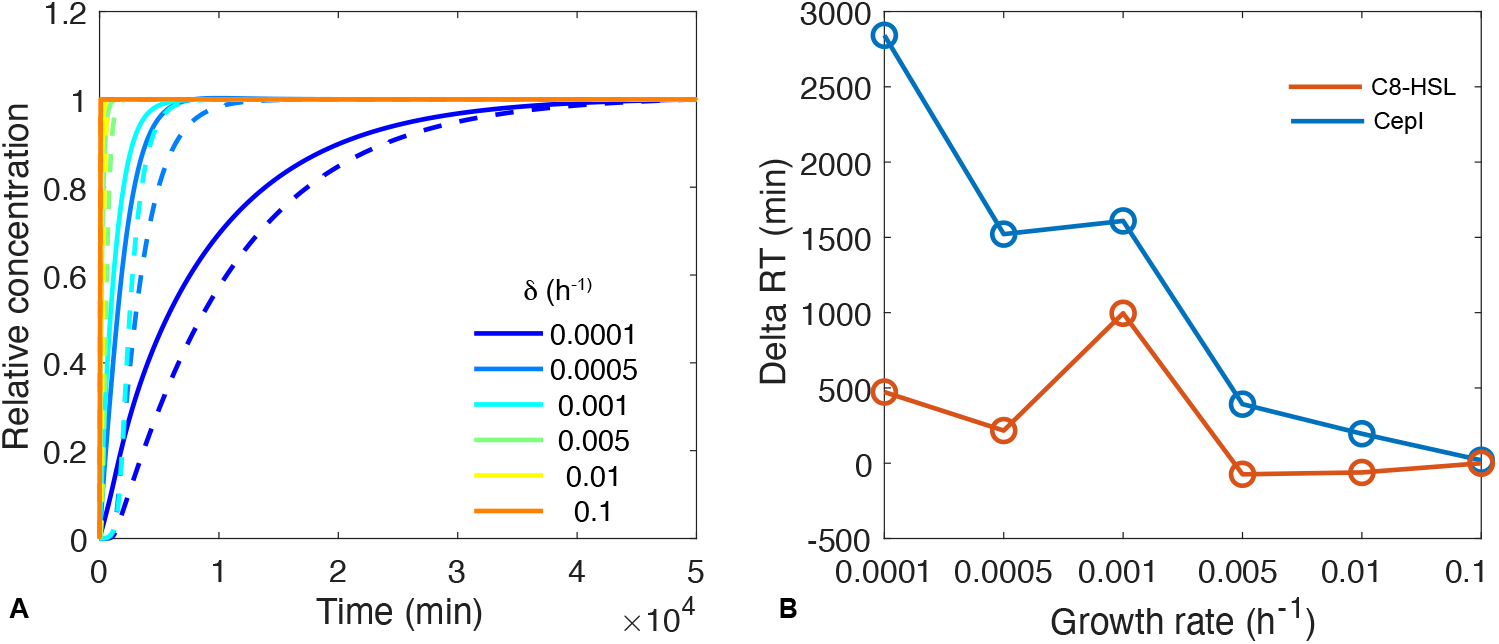
Relationship between RTs and *δ* values in the *complete / core* models. A) Dynamics of CepI concentration for different values of *δ* in the core (dashed) and complete (solid) models. B) The difference between the response time (RT) of the core model and that of the complete model (y-axis) for different growth rates tested (x-axis). Positive values of ΔRT indicate a faster RT of the *complete* model in respect to the *core* one and *vice-versa*.

**Figure 8:**
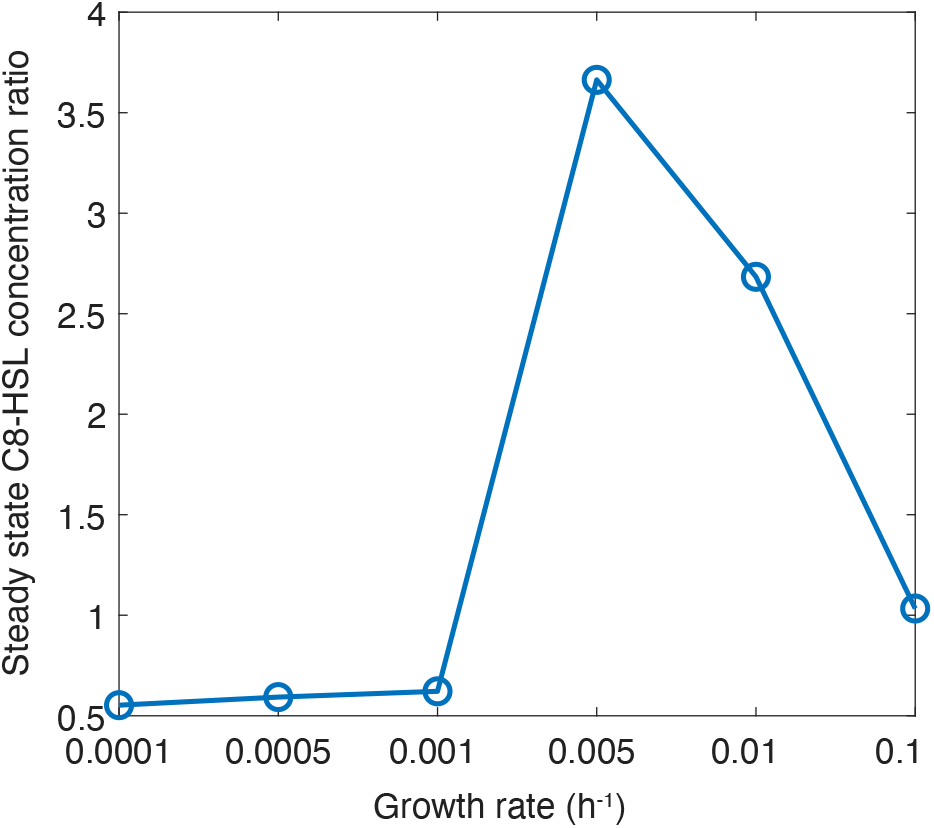
Ratio between the steady state concentration of C8HSL in the *complete* and *core* models. Higher values on the y-axis indicate higher concentrations of C8HSL at the steady state.

### 3.4. The complete architecture optimizes QS response to external AI concentration

We then evaluated the response curves of CepI, CepR and CepR* in relation to different initial AI concentration in both models. Since the external AI is produced by other bacteria from the same species, its concentration (*C8HSL^E^* in the model) can be considered a proxy of the concentration of other bacteria in the surrounding environment (i.e. cell density). We thus repeated the simulations under a wide range of initial external C8HSL concentrations (from 5 to 50 nM) and monitored the dynamics of CepI, CepR and CepR*. The results of this analysis are shown in Figure 9. With the exception of CepR, the other two proteins (CepI and the induced form of CepR, CepR*) exhibit a different behaviour in the *complete* vs. the *core* model. While in the *complete* model external concentration of AI does not influence the dynamics of CepI to its steady state level, in the *core* model the way CepI reaches its steady state concentration is strongly dependent on the external AI concentration. With higher concentrations of AI, in particular, CepI concentration, soon after its activation, is predicted to reach levels that are very far from the optimal, steady state concentration. The same trend is observed for CepR*, albeit with a much smaller discrepancy between the two architectures (Figure 9). In the *core* and *complete* schemes, CepR* reaches a concentration that is up to 8 and 3 times higher than the final steady state concentration, respectively. This difference in the reponse of the two architectures becomes even more evident by plotting the response dynamics of the 3 main species of the models (CepI, CepR and CepR*) in a two-dimensional (CepI vs CepR and CepI vs CepR*) phase space. Figure 10 shows the dynamical trajectory from an uninduced steady state to an induced equilibrium (with different initial concentrations of C8HSL, from 0 nM to 50 nM). Eventually, all the simulations ends up in their respective steady state but the trajectories are very different in the case of the *core* and *complete* models, with the first architecture leading to much higher fluctuations of CepI-CepR and CepI-CepR* systems from their optimal steady state concentrations. This is particularly evident in the CepI-CepR* phase plot where the two species are shown to steadly and gradually reach their steady state, regardless of the external initial concentration of the inducer. Taken together these results indicate an optimized control of QS response (and explicitly of CepI) in the strains harbouring the complete architecture in relation to the external AI concentration. In particular, the complete configuration seems to be able to buffer the huge spike in CepI expression level observed in the *core* model upon its activation.

**Figure 9:**
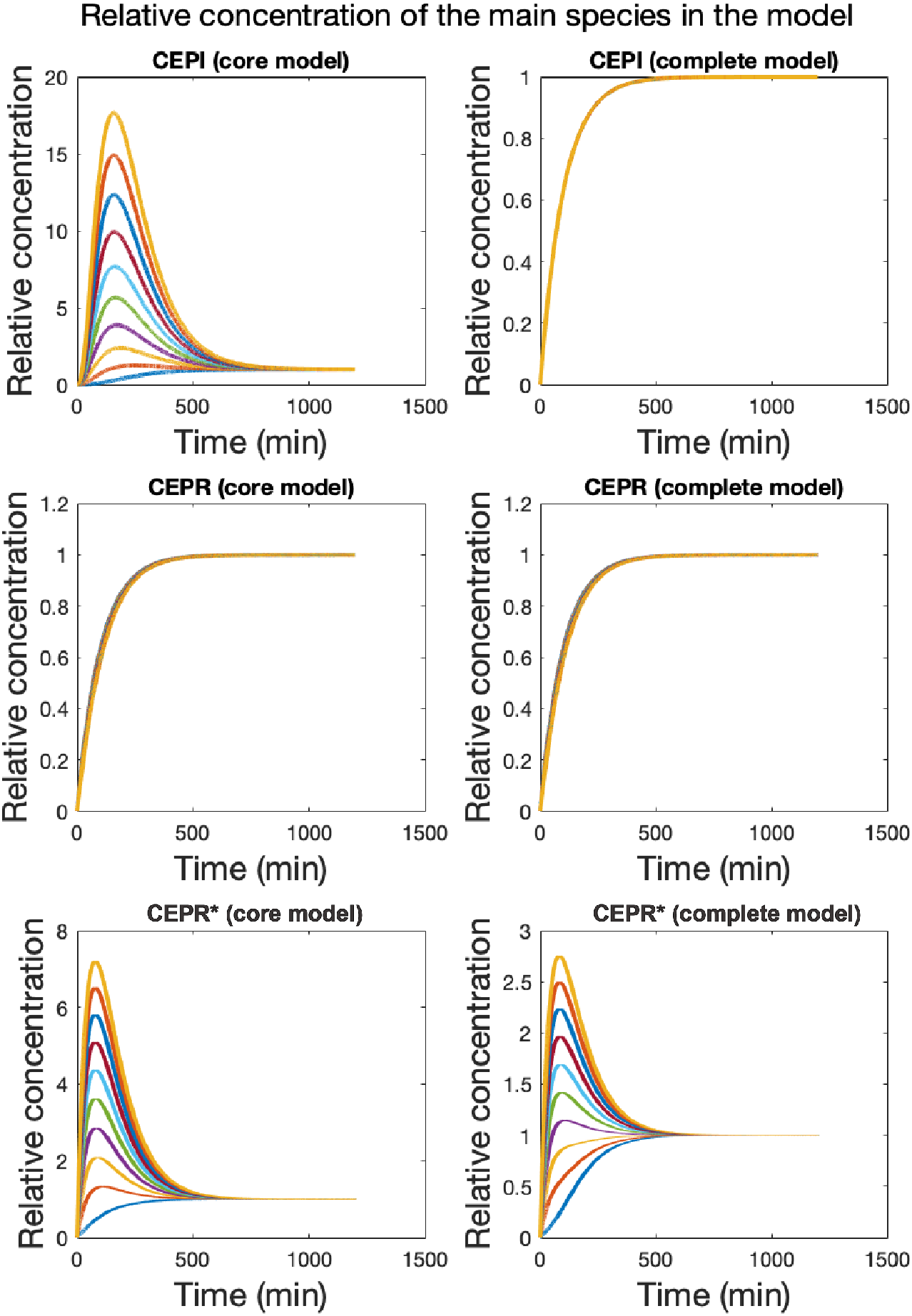
Dynamics of the main species in the model in relation to different initial concentrations of C8HSL

**Figure 10:**
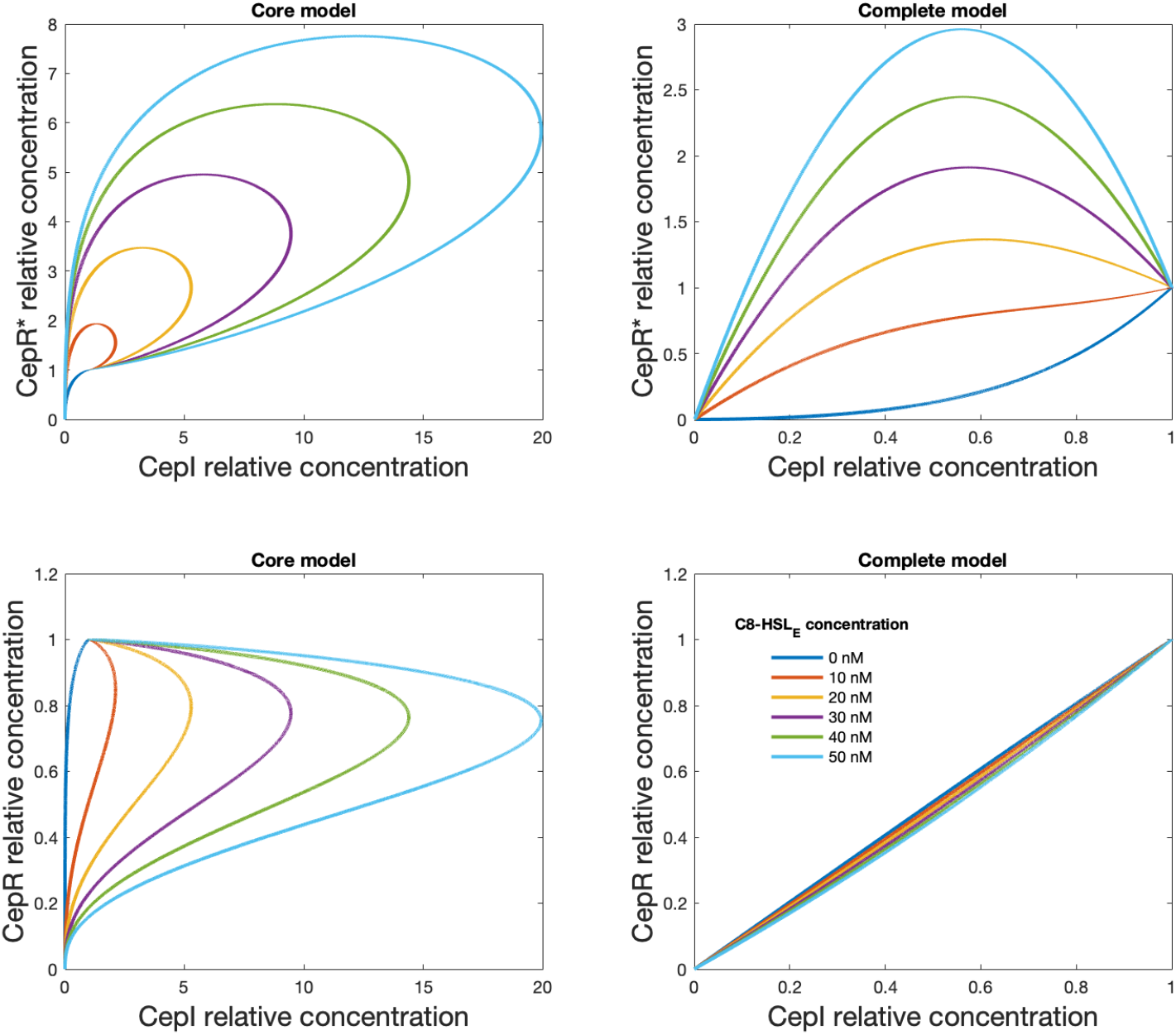
Two-dimensional phase space of CepI, CepR and CepR* dynamics in the *core* and in the *complete* models. For each species the relative concentration is plotted in each phase space, i.e. the ratio between the instantanueous and the steady-state concentrations.

### 3.5. The core model is less sensitive to stochastic fluctuations

Protein translation is a complex mechanism that is known to be affected by two different kind of noise, intrinsic and extrinsic noise. While extrinsic noise is essentially related to the environment, intrinsic noise is caused by stochastic fluctuations arising from gene expression [19, 39]. This latter aspect becomes particularly relevant when the number of proteins present in the system is relatively small, causing large deviations from the mean value of the final protein product. To overcome such problems, biological systems have developed different elementary regulatory functions that confer the system the ability to control noise. The non-coding RNAs and more specifically the microRNA-mediated feedforward loop [35, 15] are representative examples of biological networks that perform noise buffering. Similarly, QS regulation have been shown to implement regulatory loops that lead to information benefit through dedicated systems to control gene expression noise, and improving the signal-to-noise ratio [44, 28]. Thus the question one may ask is which is the role of the additional QS control system with respect tho the size of stochastic fluctuations?

To answer this question we have simulated the stochastic model described by equations (1)-(24) and we have monitored the mean (*μ*) and the standard deviation (*σ*) of two key molecular species, *CEPR*\* and *CEPI*, varying the dilution rate *δ*. Figure 11 summarizes the outcome of such analysis. As it can be seen from the plots of the first column of Figure 11, for *delta* ⩾ 3 · 10^−3^, the presence of the complete model deeply modifies the average stationary state of the concentrations of *CEPI*, increasing by a factor up to 250 the corresponding mean value of the core model. This effect is less marked for *CEPR*\*. Since the two models exhibited quite different stationary states, we chose to measure the noise strength through the standard deviation instead of the more frequently used coefficient of variation. The inspection of *σ*, indeed, reveals that fluctuations about the steady state are much more pronounced in the complete model, while the stochastic trajectories of the core model appear less noisy. This effect can be better visualized in Figure 12 where we plotted one realization of the two stochastic models for a specific values of *δ*.

**Figure 11:**
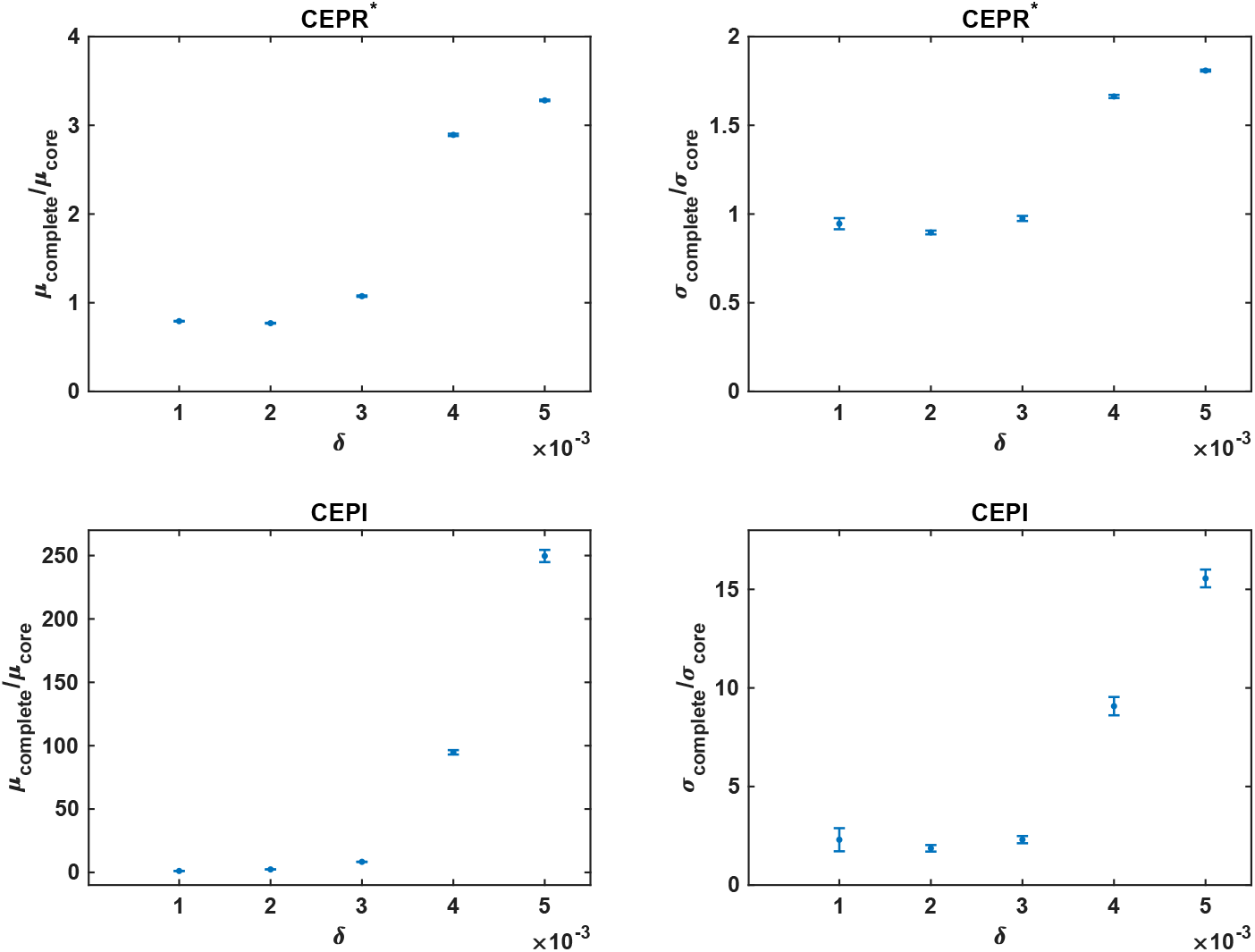
Noise properties of the complete model vs the core analogous. Top panels refer to species *CEPR*\* while bottom panels to species *CEPI*. The left plots show the behavior of the ratio of the mean of the complete model to the mean of the core model as a function of *δ*. Panels on the right show the ratio of the standard deviation of fluctuations. In all plots each point represents the value obtained averaging over 20 independent stochastic realizations of the Gillespie algorithm [13]. Error bars denote the standard deviation. Parameters values are listed in 1 while *V* = 500.

**Figure 12:**
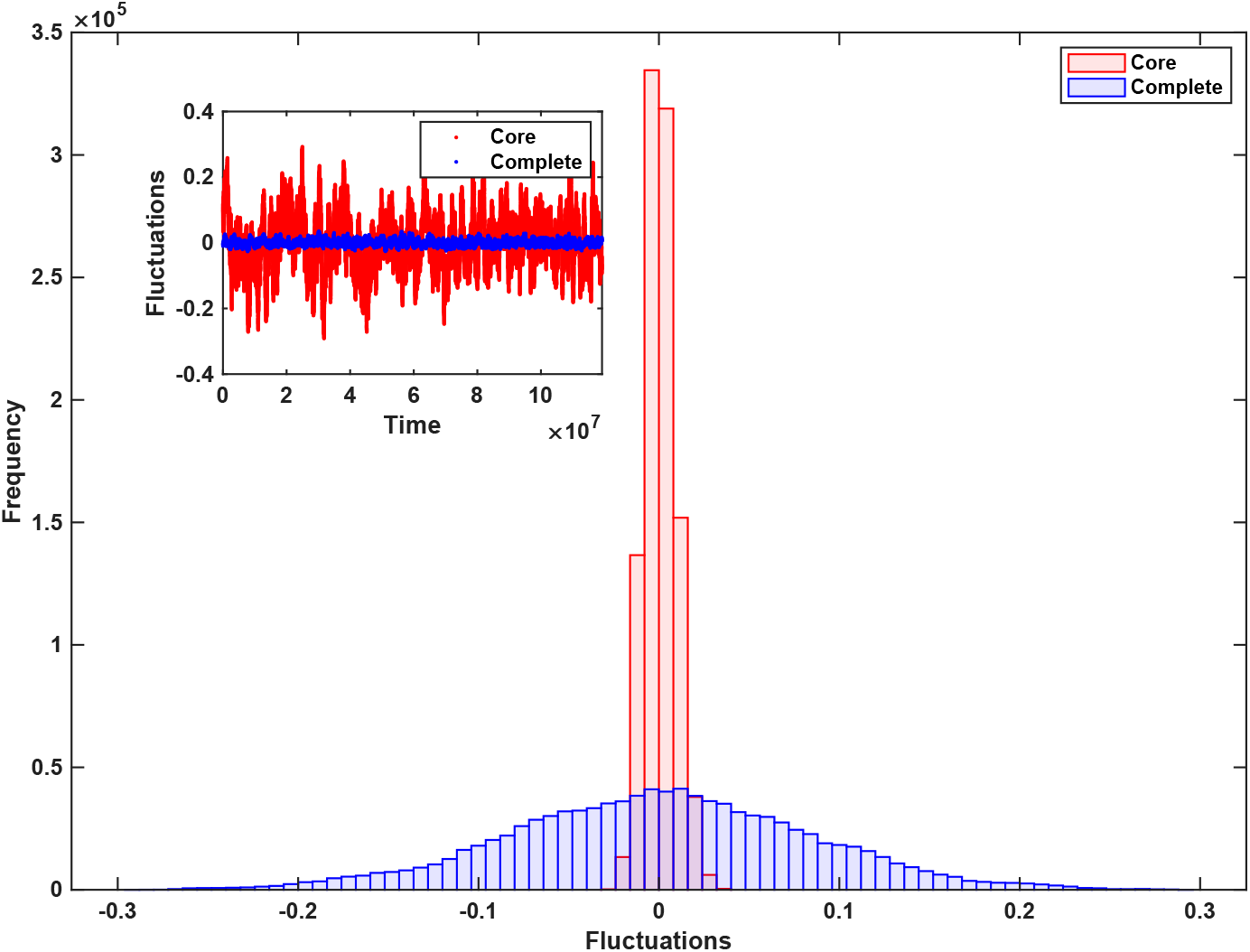
The main panel shows the comparison between the fluctuations about the steady state of *CEPI* for the complete (blu bars) and the core (red bars) model. The inset reports the corresponding single stochastic realization implemented through the Gillespie algorithm [13]. The dilution parameter *δ* is set equal to 4 × 10^−3^ while the other values are listed in Tab. 1. The volume of the system is *V* = 500.

Overall, it appears that the steady-state concentration of the species in the *core* model is less affected by stochastic fluctuations. On the contrary, the *complete* model appears to be more sensitive to random changes in the concentration of the different species, reflected by larger fluctuations around their steady state values. At the population level, the *core* organization of QS regulation might lead to a homogeneous population in which the concentration of all the species included in the model is highly similar in all the cells. Conversely, a population composed of strains harbouring the complete architecture would result heterogeneous, with all the nodel species assuming a (stochastically) different concentration around their steady state value.

## 4. Discussion

In this work, we have analysed the distribution of the *cci* genomic island in the genus *Burkholderia* and, in particular, of the QS regulatory system *cciIR* that is associated to this DNA region. This analysis pointed to a complex evolutionary history of the *cciIR* system, mainly driven by gene gain/loss events. This led to the presence of two alternative configurations of QS regulation in this group of bacteria, one (*core*) represented by the canonical and well-known CepIR regulatory scheme (homologous to the model LuxIR *Vibrio* system) and a more interconnected one (here named *complete*) in which the CepIR and CciIR systems are integrated and reciprocally influence their expression. This finding prompted us to develop a mathematical model embedding the details of both these regulatory systems and capable of addressing the following question: which are the main differences and and the possible advantages provided by the two architectures?

Overall, the simulations indicated that the main advantage of the *complete* vs. the *core* configuration may reside in a reduced response time and in an optimization of the QS response in relation to the external concentration of AI. For the *Burkholderia* representatives possessing the complete configuration, our analysis estimated a CepI response time (i.e. the time required to reach its half-maximal concentration) of 69 minutes, about 200 minutes shorter than the one of the strains with the *core* configuration of QS regulation. Interestingly, the response time appears to be linked to cellular growth rate (*δ*), with slower values of *δ* enhancing the discrepancy between the two configurations (i.e. the *complete* being faster than the *core*). Besides CepI, also the dynamics of the AI (C8HSL) appears to follow the same trend. The regulation of QS and QS-related processes has been almost exclusively interpreted as (cell) density-dependent feature. However, mounting evidence suggests that growth rate, in addition to cellular concentration, plays a major role in triggering this complex phenotypic response [31]. In *V. fischeri*, for example, nutrient deficiency (and hence reduced growth rates) promotes AI activity [45]. Similarly, it was recently shown that in *B. glumae*, there exists a growth rate threshold below which QS engages in the regulation of rhamnolipids [31]. Finally, in *P. aeruginosa*, slow growth was identified as a key condition inducing regulation of QS-controlled genes [29]. Our results confirm this link between the evolution of QS regulation and also suggest that the advantage provided by the *complete* configuration (i.e. shorter response time of QS regulation) is maximal at lower growth rates. Growth rate is known to be an important factor in microbial pathogenicity and in the outcome of infections in general [42, 6]. In real infections bacteria usually face oxygen-limited/nutrient-starved conditions and this negatively impacts cellular growth rate. Accordingly, the acquisition of the alternative feedback loops provided by the *cciIR* system, might ensure a prompt and efficient response during bacterial infections. As a matter of fact, the activity of QS-regulated enzymes (especially proteases) has been shown to be negatively correlated to growth rate in conditions that mimick the ones encountered during the infection of CF patients’ lungs [27]. Also, in a survey of Bcc strains, those strains with signs of being adapted to chronic infections (i.e. higher MICs and resistant to many antimicrobial classes) showed overall reduced growth rates [37].

Besides growth rate, we also found that the surrounding cell density may magnify the difference in QS regulation of the two circuits considered here. Indeed, we found that the path towards the optimal, physiological expression level of CepI is not influenced by the external concentration of the AI (an approximation of cell density) in the *complete* model. On the contrary, this is strongly affected in the *core* model. If one considers the steady-state level of a protein as its optimal cellular concentration and “the ultimate design goal” of an evolved regulatory circuit [40], then the advantage provided by the integrated CepIR-CciIR regulatory system may reside in allowing the cell to rapidly and efficiently reach this physiological CepI concentration, regardless of the external conditions. Taken together, the results of the simulations tests on the response of the two circuits in different physiological and environmental conditions revealed that the efficacy of the *complete* model over the *core* one is maximized in two conditions: i) low growth rates and ii) high cell density. It is thus important to consider when eventually bacteria may encounter these two circumstances. As suggested by [31], low growth rates are typically encountered by free living bacteria exposed to nutritional stress during the establishment of host infection (restricted nutritional conditions with low cell density). Additionally, combined low growth rates and high cell density are encountered by bacteria that live in biofilms inside their host. Indeed, it is known that during bacterial infections, including those of the CF lung, iron and oxygen are often limiting and this constrains the growth of pathogens to very low doubling time values [27]. In this context, it is interesting to notice that 7 out of 8 *B. cenocepacia* strains harbouring the complete architecture have been found in association with host infections (6 involving humans and one involving plants), according to the recent classification proposed in [47].

When examining the behaviour in response to stochasticity, we again found a profound difference in the two architectures. While the *core* scheme displayed limited fluctuations around the steady state, a larger variability was found for the *complete* architecture. This, in our opinion, points towards the possibility that the two configurations may give rise to homogenous vs heterogeneous populations of bacteria implementing the *core* and the *complete* circuits, respectively. It is straightforward to ask in which condition one of the two circuits may provide possible selective advantages over the other. On one side it is easier to interpret the evidence that most of the species of the Bcc do not carry the complete circuit as the result of an optimization process that gives the organisms an efficient way to control and reduce fluctuations. Nonetheless, higher levels of noise are not always synonym of adverse conditions. A more dynamic level of protein may confer the bacteria the necessary readiness to quickly react when stressed by external perturbation, thus increasing their adaptability. In recent years the role of noise in biological systems has been interpreted in different functional ways [10]. Drafting a parallel with the theory of facilitated variation [12], for example, the complete QS system seems to generate not only two output states (QS completely on or off) but, rather, a large, random array of them where most of the cells have different internal concentration of the species represented by the model, likely at a greater energy expense compared to the *core* one. In a successive selection step, the onset of a particularly advantageous condition stabilize a small fraction of these states (i.e. those cells possessing an expression pattern of QS regulated genes compatible with such a novel condition) that will proceed in the onset of the specific phenotype. In this sense, one of the possible advantage of a noisy regulatory scheme might reside in enabling the random differentiation of bacterial cells that, otherwise, would result identical. This phenotype may ultimately result in the a bet-hedging strategy in which the forthcoming effects of external perturbations are anticipated by a (random) fraction of the entire population that, on the other hand, suffers a fitness defect in normal (constant) condition [23]. Microbial species that, like the representatives of the Bcc and, especially those harbouring the *complete* QS regulation system, are capable of thriving in many disparate conditions and give rise to different phenotypes (e.g. biofilm during human infections) may benefit from such a bet-hedging strategy when facing life-style switches induced by modified environmental conditions (e.g. host invasion). If we interpret the activation of the QS-regulated gene expression program as a commitment point in cellular life (similar to the onset of other processes such as cell duplication or sporulation) cell-to-cell variability might result in significant changes in its timing from cell to cell. This would probably be due to the rates at which the master regulator of the cellular process (CepR* in our case) accumulate in the cytoplasms of the different cells. Ultimately, this may result in an broad distribution of commitment stating points inside the population, with cells expressing a wide array of genes before/after the population “mean” commitment starting point (procrastination/anticipation of differentiation).

In this context, the observation that the readiness to quickly react to (or even anticipate) the drawbacks of external perturbations provided by the *complete* architecture is in perfect agreement with the faster response time of the complete model that we have discussed in the previous section.

Based on these considerations, we speculate that the additional feedback loops provided by the CciIR regulation system may confer an evolutionary advantage to the harbouring strains during the onset and maintenance of host infections and, more in general, with their eclectic and plastic lifestyles.

## 5. Conclusions

In this work we have combined comparative genomics with mathematical modelling to elucidate the possible evolutionary advantages provided by the implementation of extra control loops in the regulation of the most important bacterial communication system, namely quorum sensing. Our study represents the first mathematical model of QS in *Burkholderia*. While the evolutionary history that led to the current configuration of QS regulation in this genus deserves further inspection, our approach suggests that additional feedback loops encoded by the CciIR system may be crucial in the optimization of QS regulation during host invasion, thus highlighting once more the importance of this process in the establishment and maintenance of bacterial infections. Work to experimentally verify model predictions is currently in progress.

## 6. Methods

### 6.1. Distribution of cci island encoded-genes in Burholderia

The entire set of *Burkholderia* complete genomes available at May 2020 was downloaded from NCBI. A total of 289 genomes were retrieved. The complete list of all the replicons and relative accession codes used in this work is available at *https://github.com/mfondi/QS_modelling*. Each gene encoded by the reference *B. cenocepacia* J2315 *cci* island was then used as query to probe this entire set of genomes retrieved in order to map the distribution of all the *cci*-encoded genes in *Burkholderia*. The Bidirectional Best Hit (BBH) approach was used to this aim, setting an E-value threshold of 1 * *e*^−20^ during BLAST [2] searches. The resulting presence/absence matrix was visualized in the form of a heatmap using R [38]. Further, in order to group the *Burkholderia* genomes on the basis of their genomic relatedness, we computed the average nucleotide identity (ANI, [18]) for each pair of genomes in the dataset. The resulting matrix was used to cluster the genomes according to the shared values and to visualize the outcome in the form of dendrogram using the R built-in *hclust* clustering function. At this point, the rows of the *cci* genes presence/absence matrix were then re-ordered based on the order of the genomes in the ANI dentrogram to better visualize the relationship between genomic relatedness and cci-genes distribution.

### 6.2. Model simulations

Most of the simulations were performed using MATLAB 2019a. The built-in ODE solver *ode23* was used to solve the ODEs of the model. Stochastic simulations based on the Gillespie algorithm were implemented using C language. The codes used to perform the simulations are available at *https://github.com/mfondi/QS_modelling*.

### 6.3. Local sensitivity analysis

We used local sensitivity analysis to assess the impact of the local variation of the model parameters on the value of each of the output variables. Specifically, the local sensitivity of the solution to a parameter is defined by how much the solution would change by changes in the parameter. Here we followed the approach described in [7] and computed the local sensitivity of each variable (*x_i_*) with respect to each parameter (*p_j_*) using the *SENS_SYS* third-party MATLAB function (Garcia Molla, 2019) and expressed as:;

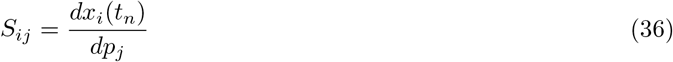

