## Supplementary Information for "The acquisition of additional feedback loops may optimize and speed up the response of quorum sensing"

*Preprint submitted to Elsevier* *June 10, 2021*

### 1. The *core* architecture of the model

From the *complete* model described in the main text, we derived the following *core* model, that matches the overall features of the one described in the main text.

$$\emptyset \xrightarrow{\alpha_{OACP}} OACP \tag{1}$$

$$CEPI \xrightarrow{\delta} \emptyset \tag{2}$$

$$CEPR \xrightarrow{\delta} \emptyset \tag{3}$$

$$CEPR^{*} \xrightarrow{\delta} \emptyset \tag{4}$$

$$OACP \xrightarrow{\delta} \emptyset \tag{5}$$

$$C8^{I} \xrightarrow{\delta} \emptyset \tag{6}$$

$$CEPR + C8^{I} \xleftrightarrow[k_{off,CEPR^{*}}]{k_{on,CEPR^{*}}} CEPR^{*} \tag{7}$$

$$OACP + CEPI \xrightarrow{\frac{V_{max,CEPI} \cdot OACP}{k_{m,CEPI} + OACP}} C8^{I} + CEPI \tag{8}$$

$$\emptyset \xrightarrow{\frac{\beta_1 \cdot (CEPR^{*})^{w}}{k_1 + (CEPR^{*})^{w}}} CEPI \tag{9}$$

$$\emptyset \xrightarrow{\frac{\beta_1}{1 + \left(\frac{CEPR^{*}}{k_3}\right)^{n}}} CEPR \tag{10}$$

$$C8^{I} \xleftrightarrow{\alpha_{C8}} C8^{E} \tag{11}$$

$$C8^{E} \xrightarrow{\delta} \emptyset \tag{12}$$

$$\emptyset \xrightarrow{\gamma_{C8}} C8^{E} \tag{13}$$

From this set of chemical equations we derive the following set of ordinary differential equations, describing the change in time of the concentration of each species.

$$\dot{CEPI} = -\delta \cdot CEPI + \frac{\beta_1 \cdot (CEPR^*)^w}{k_1 + (CEPR^*)^w} \tag{14}$$

$$\dot{OACP} = \alpha_{OACP} - \delta \cdot OACP - \frac{V_{max,CEPI} \cdot OACP}{k_{m,CEPI} + OACP}(OACP \cdot CEPI) \tag{15}$$

$$\begin{aligned} \dot{CEPR} = & -\delta \cdot CEPR - k_{on,CEPR^*}(CEPR \cdot C8^I) + \\ & + k_{off,CEPR^*} \cdot CEPR^* + \frac{\beta_1}{1 + \left(\frac{CEPR^*}{k_3}\right)^n} \end{aligned} \tag{16}$$

$$\dot{C8^E} = -\delta \cdot C8^E + \alpha_{C8^E} \cdot C8^I + \gamma_{C8,colony} - \alpha_{C8^E} \cdot C8^E \tag{17}$$

$$\dot{CEPR^*} = +k_{on,CEPR^*}(CEPR \cdot C8^I) - k_{off,CEPR^*} \cdot CEPR^* - \delta \cdot CEPR^* \tag{18}$$

$$\begin{aligned} \dot{C8^I} = & -\delta \cdot C8^I - k_{on,CEPR^*}(CEPR \cdot C8^I) + k_{off,CEPR^*} \cdot (CEPR^*) + \\ & + \frac{V_{max,CEPI} \cdot OACP}{k_{m,CEPI} + OACP}(OACP \cdot CEPI) - \alpha_{C8^E} \cdot C8^I + \alpha_{C8^E} \cdot C8^E \end{aligned} \tag{19}$$

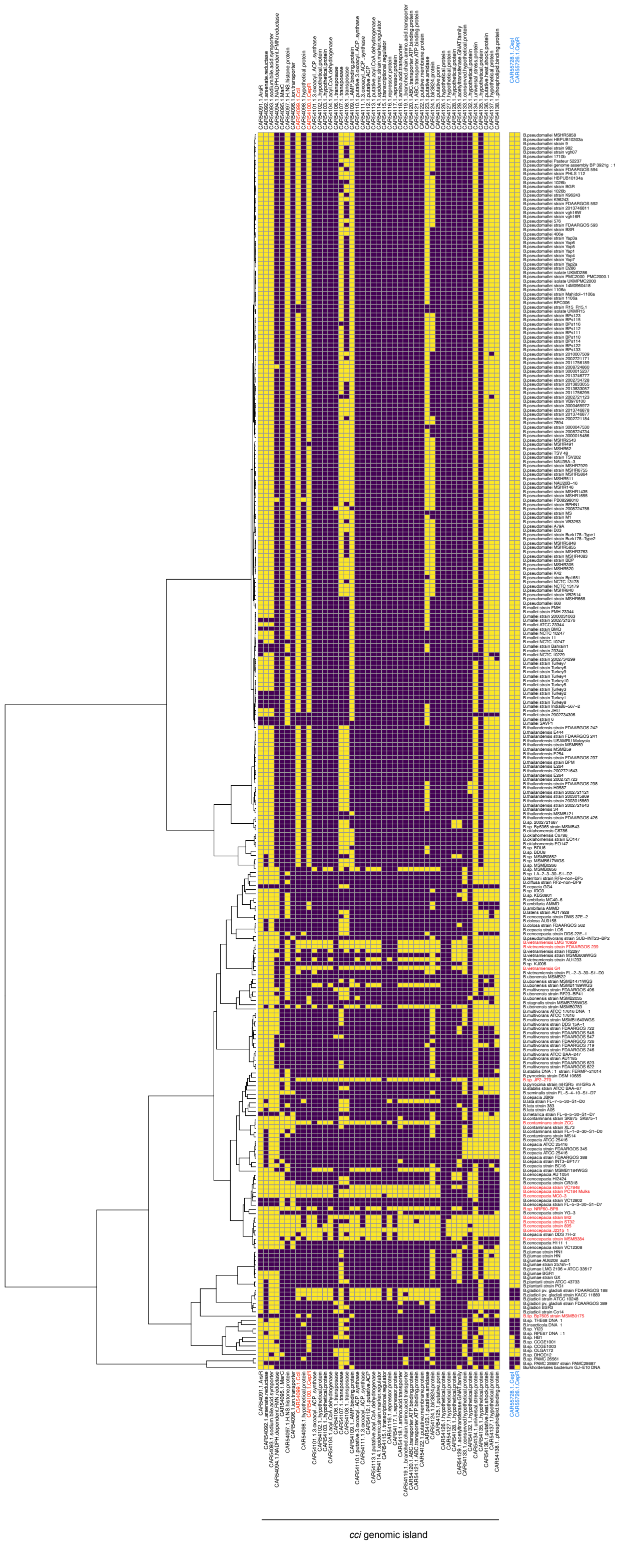


Figure S1: Distribution of *cci* island encoded genes in the entire set of complete genomes included in this work
